# Associated valence impacts early visual processing of letter strings: Evidence from ERPs in a cross-modal learning paradigm

**DOI:** 10.1101/234864

**Authors:** Mareike Bayer, Annika Graß, Annekathrin Schacht

## Abstract

Emotion effects in event-related potentials (ERPs) during reading have been observed at very short latencies of around 100 to 200 ms after word onset. The nature of these effects remains a matter of debate: First, it is possible that they reflect semantic access, which might thus occur much faster than proposed by most reading models. Second, it is possible that associative learning of a word’s shape might contribute to the emergence of emotion effects during visual processing. The present study addressed this question by employing an associative learning paradigm on pronounceable letter strings (pseudowords). In a learning session, letter strings were associated with positive, neutral or negative valence by means of monetary gain, loss or zero-outcome. Crucially, half of the stimuli were learned in the visual modality, while the other half was presented acoustically, allowing for experimental separation of associated valence and physical percept. In a test session one or two days later, acquired letter string were presented in an old/new decision task while we recorded event-related potentials. Behavioural data showed an advantage for gain-associated stimuli both during learning and in the delayed old/new task. Early emotion effects in ERPs were limited to visually acquired letter strings, but absent for acoustically acquired letter strings. These results imply that associative learning of a word’s visual features might play an important role in the emergence of emotion effects at the stage of perceptual processing.

## Introduction

Emotional valence has been shown to impact visual perception, directing attention in service of preferential processing (Pourtois, Schettino, & Vuilleumier, 2013; Vuilleumier, 2015). In event-related potentials (ERPs), this preferential processing of emotional stimuli is visible already at the stage of perceptual encoding: Emotional content increases activation in the extrastriate visual cortex, resulting in modulations of the P1 component as demonstrated for affective pictures (e.g., Delplanque, Lavoie, Hot, Silvert, & Sequeira, 2004) and facial expressions of emotion (Hammerschmidt, Sennhenn-Reulen, & Schacht, 2017; Rellecke, Palazova, Sommer, & Schacht, 2011; Rellecke, Sommer, & Schacht, 2012). Modulations in the time range of the P1, that is around 100-150 ms, have also been reported for written words (Bayer, Sommer, & Schacht, 2012; Hofmann, Kuchinke, Tamm, Võ, & Jacobs, 2009; Keuper et al., 2013, 2014; Kuchinke, Krause, Fritsch, & Briesemeister, 2014; Rellecke et al., 2011; Scott, O’Donnell, Leuthold, & Sereno, 2009).

In the case of written language, emotion effects within 200 ms after stimulus onset deserve special attention: Pictorial stimuli, including affective pictures (e.g., of a spider) or emotional facial expressions, convey their emotional content by means of their physical shape. Increased attention towards these stimulus features has often been related to biological preparedness (Öhman & Mineka, 2001) and survival-relevance (Lang & Bradley, 2010). Written words, however, consist of arbitrary symbols, which require the translation into meaningful contexts. Most reading models assume that initial orthographic analyses takes around 200 ms, and is only then followed by the processing of lexico-semantic features, as, for example, visible in the N400 component of ERPs (for review, see Barber & Kutas, 2007). In line with these assumptions, previous research has localized so-called lexicality effect, i.e. the difference in ERPs between existing words and (orthographically legal) pseudowords, at around 200 to 400 ms after word onset (Palazova, Mantwill, Sommer, & Schacht, 2011; Schacht & Sommer, 2009; but see Sereno, Rayner, & Posner, 1998 for earlier effects of lexicality). Importantly, these studies consistently reported effects of emotional content to coincide with or even to follow the ERP differences between legal words and pseudowords. In the light of these findings and accounts, emotion effects at approximately 100 ms after word onset for written words are remarkable and seem to allow for two possible explanations: First, it might be possible that lexico-semantic access occurs faster than assumed by serial reading models, and that later ERP effects (like the N400) reflect recurrent processing instead of initial lexico-semantic activations. Evidence for this assumption is provided by a number of studies showing effects of (non-emotional) lexico-semantic variables within 200 ms after stimulus onset, including word frequency, but also semantic knowledge (Hauk, Pulvermüller, Ford, Marslen-Wilson, & Davis, 2009; Pulvermüller, Shtyrov, & Hauk, 2009; Rabovsky, Sommer, & Abdel Rahman, 2012; Sereno & Rayner, 2003).

Second, it seems possible that early emotion effects in response to written words do not reflect lexico-semantic processing, but are rather based on associative learning of stimulus valence, which might become tagged to a word’s shape. There is accumulating evidence outside of the language domain suggesting that associative learning of perceptual features can impact the early stages of visual perception: In a study by Schacht and co-workers (Schacht, Adler, Chen, Guo, & Sommer, 2012), formerly unfamiliar Chinese characters were associated with positive, neutral or negative monetary outcome by means of associative learning. In a test session 1-2 days later, characters associated with monetary gain elicited an increased posterior positivity at around 150 ms after stimulus onset. Using a similar associative learning paradigm and peripherally presented textures, Rossi and colleagues (Rossi et al., 2017) showed that associative learning of perceptual features can impact even earlier stages of stimulus processing in the primary visual cortex as indexed by the C1 component peaking at around 75 ms after stimulus onset. Finally, associative learning of stimulus valence has recently also been demonstrated for biologically relevant stimuli: Neutral faces that were previously associated with monetary reward elicited increased amplitudes of the P1 component (Hammerschmidt et al., 2017).

In addition to the studies reported above, a number of studies have employed classical conditioning paradigms in order to pair unconditioned stimuli with aversive/appetitive stimuli. Similar to the studies reported above, these investigations report rapid effects of conditioned valence starting from 50 ms after stimulus onset, both for abstract stimuli (Hintze, Junghöfer, & Bruchmann, 2014; Stolarova, Keil, & Moratti, 2006) and stimuli with biological relevance (Pizzagalli, Greischar, & Davidson, 2003; Rehbein et al., 2014, 2015). Interestingly, a number of these studies have shown that even rapid effects are not merely based on modulations in sensory areas, but also involve activations in prefrontal areas (Hintze et al., 2014; Steinberg, Bröckelmann, Rehbein, Dobel, & Junghöfer, 2013). Finally, Montoya and colleagues (1996) paired pseudowords with electric shocks; they report enhanced N100 amplitudes in a subsequent old/new recognition task for shock-associated words compared to non-shock words. In two studies using an emotional conditioning paradigm (Fritsch & Kuchinke, 2013; Kuchinke, Fritsch, & Müller, 2015), pseudowords were paired with emotional pictures and subsequently elicited effects of conditioned valence around 100 to 150 ms. Since pseudowords do not have a pre-existing meaning, these studies demonstrate that also meaningless letter strings can be tagged with conditioned valence.

Taken together, previous literature provides evidence for both fast semantic access and associative learning accounts. However, both mechanisms are not necessarily mutually exclusive, but might even interact. Crucially, there is no way of distinguishing between the two options using existing words as stimulus materials, because emotional valence and perceptual features are pre-determined and cannot be established experimentally. Therefore, the present study used meaningless letter strings and a cross-domain design in order to determine whether associated valence might become tagged to stimulus shape and elicit early ERP modulations. In a learning session, letter strings were associated with positive, negative or neutral valence by means of monetary gain, loss, or neutral outcome. They were presented again in a test session 1-2 days later, while EEG was recorded. Importantly, half of the stimuli were acquired in the visual domain, while the other half was acquired in the auditory domain. In the test session, all letter strings were presented in the visual domain. As a result, stimuli were associated to the same valence categories in both learning modalities, but participants gained experience with perceptual features only for stimuli acquired in the visual modality. Importantly, our letter strings did not acquire any sort of semantic meaning beyond these valence associations (i.e., they were not associated to a specific meaning), thus enabling an unconfounded investigation of associated valence effects. Similar procedures have been chosen in the investigation of prosodic (Paulmann & Kotz, 2008) and syntactic information (Goucha & Friederici, 2015).^1^ Rather than employing classical conditioning, in the present study we used an associative learning paradigm highly similar to previous studies (Hammerschmidt et al., 2017; Rossi et al., 2017), since these studies provided evidence for effects of associated valence both for abstract symbols and faces. Here we aimed at expanding these findings to rather complex, pseudo-linguistic stimuli, i.e. pronounceable 4-letter strings.

In addition to early effects in the P1 time range, we investigated modulations of the P300, and, in the context of emotional processing, of the LPC (Late Positive Complex). The LPC was related to increased higher-order stimulus evaluation of emotional content (Cuthbert, Schupp, Bradley, Birbaumer, & Lang, 2000). The P300, more generally, is thought to index attention allocation, memory processing and task relevance (Johnson, 1986; Polich, 2007). Concerning effects of associated valence, previous findings are inconclusive, with some studies reporting LPC modulations (Fritsch & Kuchinke, 2013; Hammerschmidt, Kagan, Kulke, & Schacht, 2018; Schacht et al., 2012), while others do not (Hammerschmidt et al., 2017; Kuchinke et al., 2015). Interestingly, Rossi and colleagues reported that LPC modulations were limited to a categorisation task, but were absent in an old/new task, being in line with the task sensitivity of the P300/LPC component (e.g., Schacht & Sommer, 2009).

In accordance with previous literature (Hammerschmidt et al., 2017; Schacht et al., 2012), we expected effects of associated valence in visual event-related potentials within the first 200 ms after stimulus onset. In case these effects would be based on fast access to associated valence information, i.e. the valence category of a letter string, these effects should occur irrespective of learning modality. If, however, associative learning would be based on a letter string’s perceptual features, early ERP effects should be limited to pseudowords acquired in the visual domain. In contrast, these effects should be absent for letter strings learned in the acoustic modality, since participants had no experience with their visual features. Concerning the direction of early effects of associated valence, previous literature is inconsistent, showing both increased amplitudes for positive associated valence (Hammerschmidt et al., 2017; Schacht et al., 2012) and for negative associated valence (Rossi et al., 2017), as well as decreased amplitudes for negative associated valence (Fritsch & Kuchinke, 2013). Therefore, we did not specify hypotheses about the direction of early effects of associated valence. Concerning the P300, we expected to replicate previous findings of increased amplitudes for associated letter strings compared to novel distractors in terms of a classical old/new effect (Kuchinke et al., 2015; Rossi et al., 2017; for overview, see Rugg & Curran, 2007), but did not make predictions on effects of associated valence, reflecting the inconsistency in previous literature.

## Methods

### Participants

Data were collected from 73 participants; eight datasets had to be discarded due to excessive EEG artefacts (5) and poor accuracy in the testing session (3). All remaining 65 participants (50 women, mean age = 24.8 years, SD = 3.7 years, 4 left-handed) were native speakers of German, had normal hearing and normal or corrected-to-normal vision.^2^ Participants were compensated with course credit or 8 Euros per hour; additionally, participants received the money they had earned during the learning session (mean final balance = 14.41 Euro, SD = 3.5 Euro, including a base pay of 3 Euro).

### Stimuli

Target stimuli consisted of 24 disyllabic letter strings following the phonological form consonant – vowel – consonant – vowel (e.g., foti, metu, bano). They were constructed in accordance with phonological rules of German and followed phoneme-grapheme correspondence. Letter strings were distributed to 3 *valence groups* of 8 words each. Within each valence group, four words were presented in their written form in the learning session, while the remaining four words were learned in the auditory domain. Analyses of control variables were referred both to valence groups and to valence x domain subgroups. All groups were controlled with regard to sublexical bigram frequency (character bigram frequency, obtained from the dlex-database accessible at www.dlexdb.de; Heister et al., 2011), all *F*s(5,18) < 1; see Table 1. For auditory presentation, stimuli were spoken by a male speaker in neutral prosody and stressed on the first syllable in order to achieve a natural German pronunciation. Acoustic control variables included word duration, mean and peak amplitude, and mean fundamental frequency (F0); all *F*s(5,18) < 1.

**Table 1.**
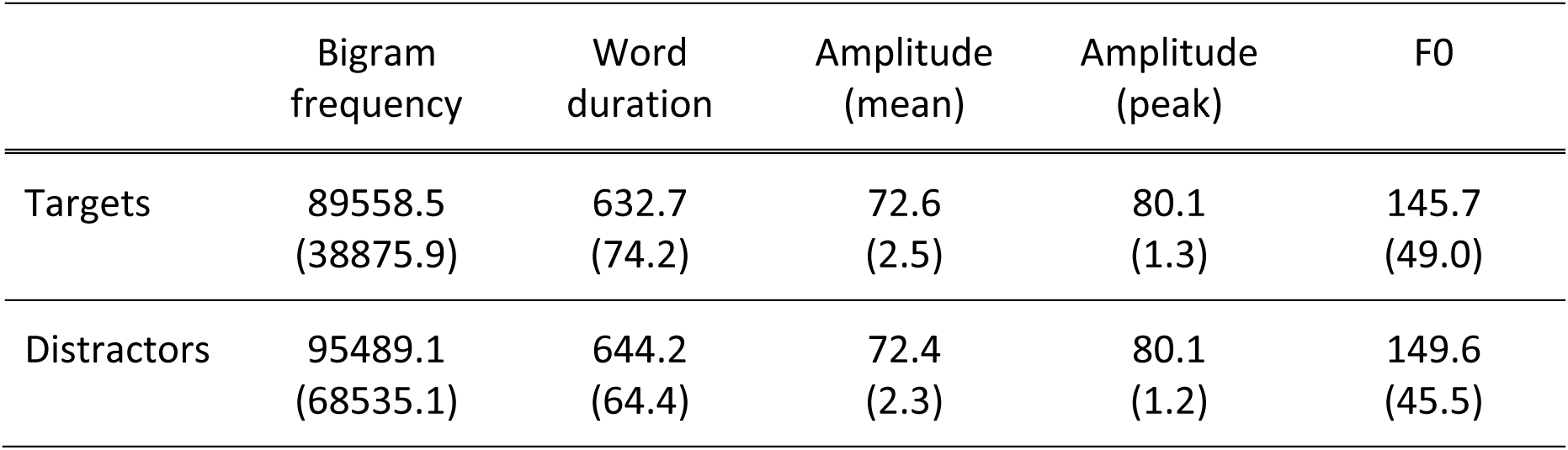
Descriptive statistics (means and standard deviations) of linguistic and auditory stimulus variables. Bigram frequency was obtained from the dlex-database. Word duration, amplitude and F0 are indicated in milliseconds, decibel and Hertz, respectively, measured with Praat (Boersma & Weenik, 2009). Amplitude values are only relevant for comparisons between experimental categories, but do not reflect actual presentation levels, which were individually adjusted to a comfortable volume level.

The assignment of valence groups and valence x domain subgroups was counterbalanced across participants in a way that each letter string was assigned to each valence category and learning modality for an equal number of participants.

For the test session, 288 distractors that followed the same structure as the target letter strings were constructed by computer algorithm, excluding orthographic neighbours of target letter strings. Analyses of all control variables listed above revealed no significant differences between targets and distractors, all *F*s(1,310) < 1.

### Procedure

The experiment consisted of two sessions. In the learning session, participants associated letter strings to monetary gain, loss, or neutral outcome by means of associative learning. In the test session 1-2 days later, all letter strings acquired in the learning session were again presented in an old/new decision task amongst unknown distractors. The study was approved by the local ethics committee of the Institute of Psychology at the University of Göttingen.

#### Learning session

Upon arrival, participants received detailed information about experimental procedure and provided informed consent as well as demographic information. A short, custom-made hearing test was conducted in order to ensure sufficient hearing.

The participants task was to acquire associations between letter strings and valence categories by pressing one of three buttons – corresponding to gain, loss, or neutral outcome – after presentation of a letter string. Since no information about the correct outcome category for each specific stimulus was provided prior to the experiment, participants had to employ a trial-and-error procedure in order to learn a letter string’s respective valence category, using the feedback that was provided after their choice. This feedback indicated the amount of money the participant had won or lost in the present trial. From this information, participants could gain two facts: First, it provided information about the valence category of a given stimulus, since gain stimuli always resulted in monetary gain, loss stimuli in monetary loss, and neutral stimuli in zero change. Second, in case of gain and loss symbols, the amount of money won or lost indicated whether participants had made a correct or incorrect classification. In case of correct classifications, participants either won more money (correct choice: +20 Cents, incorrect choice: +10) or lost less (correct: -10 Cents, incorrect: -20 Cents). For an overview of the learning scheme, see Table 2.

**Table 2:** Learning scheme. Monetary consequences resulting from correct and incorrect classifications for each outcome category.

| Valence category | Words per category | Classification | Response | Outcome |
| --- | --- | --- | --- | --- |
| Positive | 8 | Positive | Correct | +20 |
|  |  | Neutral/Negative | Incorrect | +10 |
| Neutral | 8 | Neutral | Correct | 0 |
|  |  | Positive/Negative | Incorrect | 0 |
| Negative | 8 | Negative | Correct | -10 |
|  |  | Positive/Neutral | Incorrect | -20 |

Half of the letter strings (n=12 each) was learned in the visual domain; the other half was learned in the auditory domain. The order of learning modalities was counterbalanced. In the visual learning modality, a fixation cross was shown for 0.5 s, followed by a letter string that was displayed for up to 5 s until the participant had pressed a response button. During auditory presentation, letter strings were presented via loudspeakers; the fixation cross was shown for 0.5 s prior to the letter string presentation and remained on the screen until button press for up to 5 s. After an inter-stimulus interval of 1.5 s, the feedback stimulus was presented for 1s. The feedback consisted of a light grey disk that showed the amount of money the participant had won or lost by their choice; gain symbols were presented in green, loss symbols in red, and neutral symbols in dark grey font colours, respectively. The assignment of response buttons to outcome categories (left to right: gain – neutral – loss or loss – neutral – gain) was counterbalanced.

Within each learning modality, all 12 letter strings were presented once within each block in randomized order. Between blocks, the current balance was displayed on the screen while participants were allowed a short break. For both learning modalities, the learning procedure ended after a participant had finished 30 blocks or reached a learning criterion consisting of 48 correct classifications within the last 50 responses.

#### Test session

The test session took place 1-2 days after the learning session. It consisted of an old/new decision task; participants had to indicate by button press whether a letter string had been acquired during the learning session (‘old’) or presented an unfamiliar letter string (‘new’ distractor). All stimuli were presented in the visual modality. The task consisted of 8 blocks of 48 stimuli each (24 associated letter strings and 24 novel distractors); stimuli within each block were presented in randomized order. Subjects without auditory testing session completed 12 experimental blocks. Distractors (n=192/288) were only presented once in the course of the test session. A fixation cross was presented for 0.5 s at the start of each trial, followed by 1.5 s of stimulus presentation and a 2s inter-trial interval. The assignment of buttons to response categories was counterbalanced.

### Data acquisition and preprocessing

The electroencephalogram was recorded from 64 electrodes positioned in an electrode cap (Biosemi Active Two System). Additionally, 2 external electrodes were applied to the left and right mastoids; 4 electrodes were positioned at the outer canthi and below both eyes in order to record horizontal and vertical electrooculograms. The continuous EEG was referenced online to a CMS-DRL ground (driving the average potential across all electrodes as close as possible to the amplifier zero) and recorded with a sampling rate of 512 Hz. Offline, data were re-referenced to average reference and band-pass filtered with 0.03 Hz (12dB/oct) to 40 Hz (48dB/oct); additionally, a notch filter was applied. Blinks were corrected using Surrogate Multiple Source Eye Correction (Ille, Berg, & Scherg, 2002) as implemented in BESA (Brain Electric Source Analyses, Megis Software GmbH), using default parameters for blink correction. Artefact-free, spontaneous blinks were used to obtain individual blink topographies. Electrode channels with poor signal were interpolated by 4^th^ order spherical splines (1,1% of channels on average, max. = 5 channels). Continuous data were segmented into epochs ranging from 100 ms before to 1000 after stimulus onset and referred to a 100 ms pre-stimulus baseline. Segments containing artefacts, i.e. activations exceeding ± 100 μV or voltage steps larger than 100 μV, were rejected in a semiautomatic way (3,14 % of trials on average). Furthermore, trials with erroneous responses were excluded from analyses; this resulted in differences in trial numbers per experimental condition (ranging between 86.4 and 93.6% per condition; mean = 90.0%, rm-ANOVAs show a main effect of learning modality, *F*(1,64) = 5.86, *p* < .05, and a main effect of valence, *F*(2,128) = 15.31, *p* < .001; reflecting higher trial numbers for the visual compared to the auditory modality, and for positive > negative > neutral valence). Finally, epochs were averaged per subject and experimental condition (learning modality x emotion).

### Data analyses

Learning performance was assessed as percentage of correct classifications per learning modality and valence condition. It was analysed by a repeated-measures (rm)-ANOVA including the factors learning modality (visual, auditory) and valence (positive, neutral, negative). Additionally, we calculated simple slopes of the cumulative accuracy of learning performance separately for positive, neutral and negative valence and visual and auditory learning modality and statistically analysed them using rm-ANOVA. Reaction times and accuracy rates of the test session were analysed by rm-ANOVAs with the factors learning modality (visual, auditory) and valence (positive, negative, neutral). Furthermore, we analysed reaction times and accuracy rates in response to associated letter strings (targets) and new distractors using rm-ANOVAs.

Time windows of ERP analyses were determined by visual inspection of grand mean waveforms. P1 amplitudes were analysed as main ERP amplitudes in a time window of 85 to 100 ms at occipital electrodes P7, P8, P9, P10, PO7 and PO8, corresponding to the peak latency of the P1. Amplitudes of the LPC were analysed at a group of centro-parietal electrodes (CPz, P1, Pz, P2) in the time window from 400 to 800 ms. In all analyses, the influence of learning modality and emotion was analysed with rm-ANOVAs including the factors learning modality (2), valence (3), and electrode (depending on ROI size).

In addition, we also compared ERP effects elicited by associated letter strings and new letter strings (i.e., distractors in the old/new task). Analyses – rm-ANOVAs with the factor old/new (2) and electrode (according to ROI size) – were performed on mean ERP amplitudes in the same time windows and regions of interest as comparisons of emotion categories.

Degrees of freedom in rm-ANOVAs were adjusted using Huynh-Feldt corrections. Results are reported with uncorrected degrees of freedom, but corrected *p*-values. Post-tests were performed with rm-ANOVAs and reported with uncorrected *F*-values and effect sizes, but with Bonferroni-adjusted *p*-values.

## Results

### Behavioural data

#### Learning session

The analyses of accuracy rates in the learning session showed a main effect of valence, *F*(2,128) = 34.07, *p* < .001, = *η_p_*^2^ =.347, indicating that participants classified stimuli associated with positive and negative valence with a higher accuracy than neutral stimuli, *F*(1,64) = 54.69, *p* < .001,*η_p_*^2^= .461, and *F*(1,64) = 27.10, *p* < .001, *η_p_*^2^ = .297, respectively; indexing overall facilitated acquisition. Furthermore, positive associations were acquired with a higher accuracy than negative associations, *F*(1,64) = 9.19, *p* < .05, *η_p_*^2^ = .126. There was no main effect of learning modality and no interaction between learning modality and valence, *F*s < 1. Analyses of simples slopes of cumulative accuracy (Figure 1) showed a main effect of valence, *F*(2,128) = 25.03, *p* < .001, *η_p_*^2^ = .281; based on steeper slopes for positive and negative compared to neutral stimuli, *F*s(1,64) > 23.27, *p*s < .001, *η_p_*^2^s > .267, whereas slopes between positive and negative words did not differ, F(1,64) = 4.91, p = .09. On average, participants performed 28.7 learning blocks in total (SD = 11.5).

**Figure 1.**
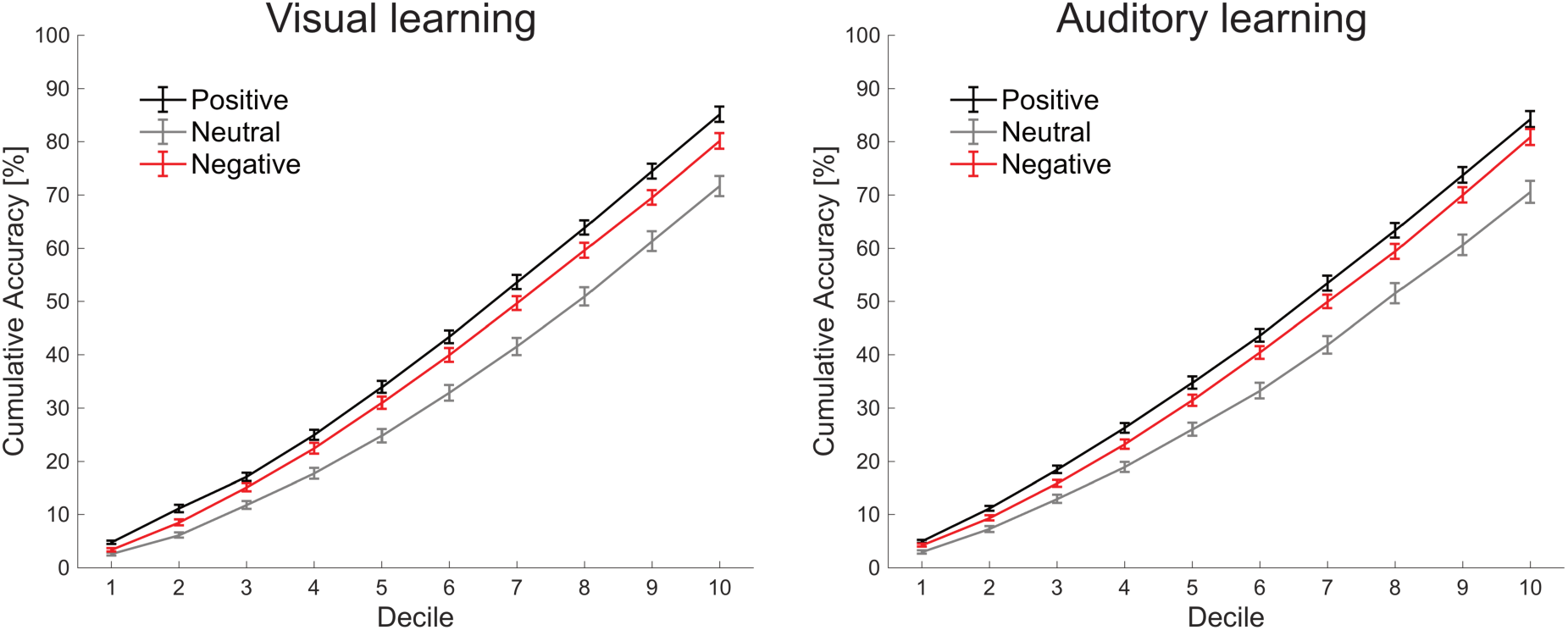
Cumulative accuracy scores per decile for visual and auditory learning sessions.

#### Test session

Behavioural results are depicted in Figure 2. Analyses of RTs revealed an effect of learning modality, *F*(1,64) = 8.86, *p* < .01, *η_p_*^2^ = .122, with faster RTs in response to letter strings acquired in the same – i.e. visual – modality than in the auditory modality. Furthermore, there was a main effect of valence, *F*(2,128) = 19.98, *p* < .001, *η_p_*^2^ = .238, due to faster RTs for positive compared to neutral stimuli, *F*(1,64) = 44.20, *p* < .001, *η_p_*^2^ = .408, and compared to negative stimuli, *F*(1,64) = 12.23, *p* < .01, *η_p_*^2^ = .160. Finally, RTs for negative letter strings were faster than in the neutral condition, *F*(1,64) = 7.01, *p* < .01, *η_p_*^2^ = .099. There was no significant interaction between learning modality and valence, *F*(2,128) = 2.049, *p* = .133.

**Figure 2.**
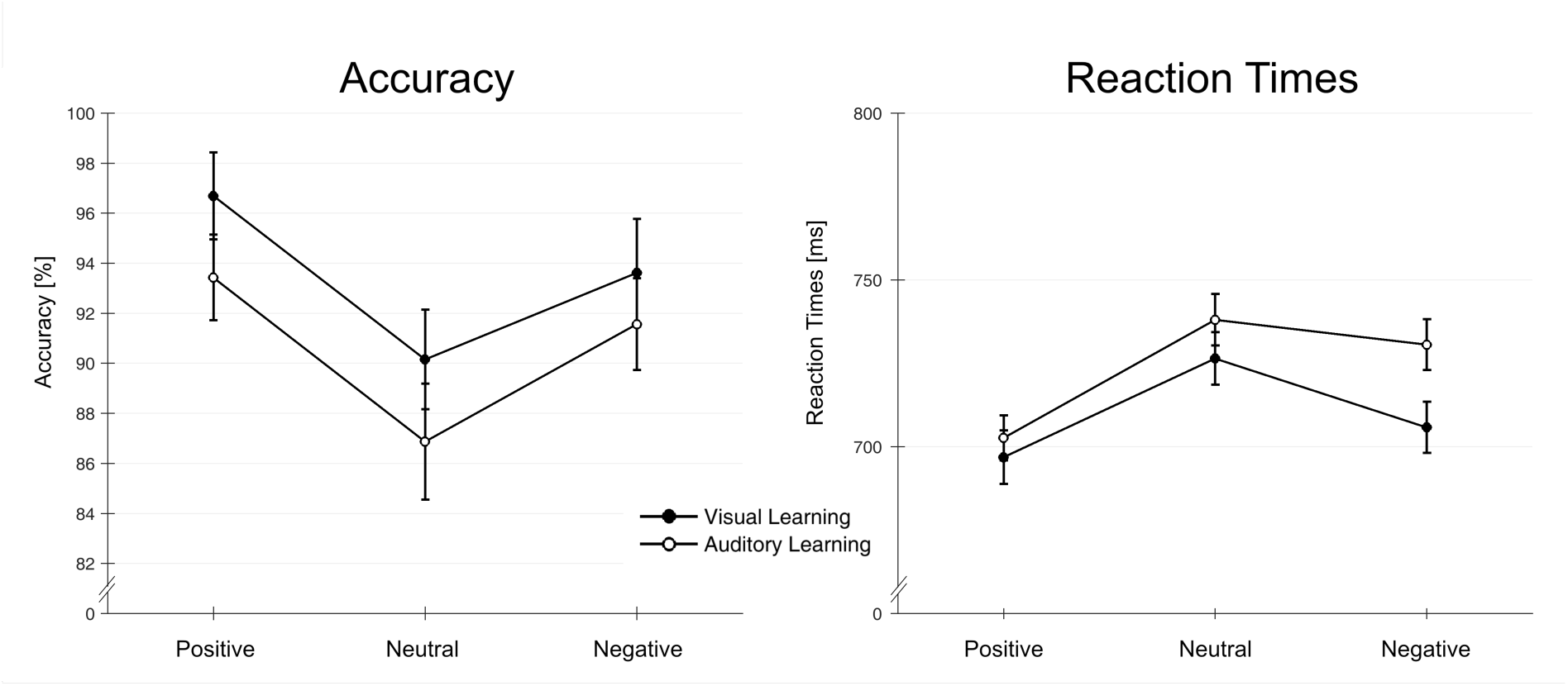
Accuracy rates and reaction times per learning modality and valence category. Error bars indicate 95% confidence intervals.

Analyses of accuracy rates showed an effect of learning modality, *F*(1,64) = 9.09, *p* < .01, *η_p_*^2^ = .124, based on higher accuracy for stimuli acquired in the visual domain. A main effect of valence, *F*(2,128) = 15.00, *p* < .001, *η_p_*^2^ = .190, reflected higher accuracy for positive and negative than neutral stimuli, *F*s(1,71) > 10.90, *p*s < .01, *η_p_*^2^s> .146. Positive and negative letter strings did not differ, *F*(1,64)= 4.73, *p* = .099, *η_p_*^2^ = .069. There was no interaction between learning modality and valence, *F*(2,128) < 1.

Accuracy rates did not show a significant difference between associated stimuli (targets) and new distractors, *F*(1,64) = 2.33, *p* = .132, but participants were significantly faster in response to targets than to distractors, *F*(1,64) = 29.16, *p* < .001, *η_p_*^2^=.313.

#### ERPs

In the time range from 85 to 100 ms, analyses of P1 amplitudes showed a trend for an interaction between learning modality and valence, *F*(2,128) = 2.73, *p* = .069, *η_p_*^2^= .041. There were no significant main effects of learning modality, *F*(1,64) = 2.26, *p* = .138, or valence, *F*(2,128) = 1.41, *p* = .248.

The analyses of LPC amplitudes between 400 and 800 ms after stimulus onset showed a main effect of learning modality, *F*(1,64) = 6.92, *p* < .05, *η_p_*^2^ = .098, reflecting higher amplitudes for letter strings acquired in the auditory domain compared to the visual domain. There was no effect of valence and no interaction of learning modality and valence, *F*s < 1.

The comparison of associated and novel letter strings (old/new effect) revealed no significant differences in the early time window from 85 to 100 ms after stimulus onset, *F*(1,64) < 1. However, P300 amplitudes between 400 and 800 ms were boosted for acquired compared to new letter strings, *F*(1,64) = 169.52, *p* < .001, *η_p_*^2^ = .726, see Figure 3C.

**Figure 3.**
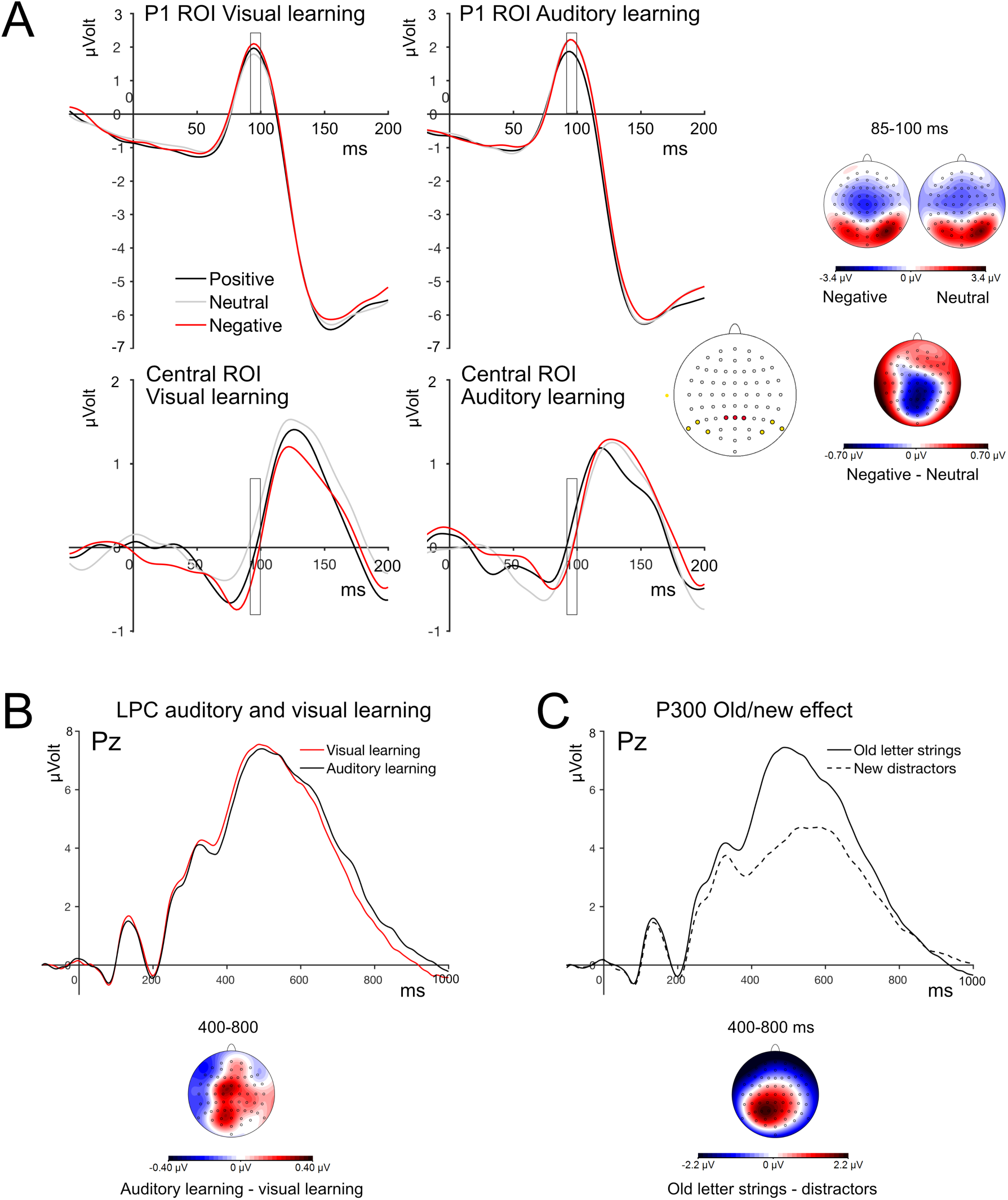
ERP results. A: P1 time window. ERP mean waveforms per valence category, separately for visually acquired letter strings (left) and acoustically acquired letter strings (right) at the P1 ROI (upper panel) and at the central ROI (lower panel). Positions of ROI electrodes are indicated in yellow (P1 ROI) and red (central ROI). Scalp topographies show distributions for neutral and negative words, and the difference topography of the two conditions. B: ERP waveforms at electrode Pz for visually and acoustically acquired stimuli. The scalp topography depicts the difference distribution for acoustically minus visually acquired letter strings between 400 and 800 ms. C. Grand mean waveforms for old letter strings and distractors at electrode Pz, and their difference topography in the P300 interval from 400 to 800 ms.

In order to confirm that the occurrence of an auditory testing session and the number of visual presentation blocks in the testing session did not influence our results, we repeated all analyses with the between-subject factor indicating session type. Results of these analyses confirmed all of our experimental findings. Importantly, they did not reveal any interaction between effects of experimental conditions and session type, all *F*s (2,126) < 1.89, *p*s > .156; and *F*s(1,63) < 1.

### Exploratory analyses

#### ERPs

Visual inspection of ERP waveforms in the P1 time window (see Figure 3) revealed a negative counterpart of occipital P1 positivities at parietal electrodes. In order to further explore the nature of early modulations, this negativity was quantified at electrodes P1, Pz and P2, in the time window from 85 to 100 ms. Analyses revealed a significant interaction between learning modality and valence, *F*(2,128) = 3.58, *p* < .05, *η_p_*^2^=.053. This interaction was based on the fact that valence effects were limited to visually acquired letter strings, *F*(2,128) = 4.05, *p* < .05, *η_p_*^2^ = .059, but were absent for letter strings acquired in the auditory domain, *F*(2,128) = 1.52, *p* = .224. Within the visual learning modality, post-tests showed an increased negativity for negative compared to neutral stimuli, *F*(1,64) = 7.95, *p* < .05, *η_p_*^2^ = .110 (see Figure 3A). No other comparison reached significance, *F*s(1,64) < 2.83, *p*s > .291. Exploratory post-tests at the lateral P1 ROI for negative vs. neutral words acquired in the visual modality corroborated this effect, showing increased P1 amplitudes for negative compared to neutral words, *F*(1,64) = 4.52, *p* < .05, *η_p_*^2^ = .066.

### Brain-behaviour correlations

Correlation analyses between EEG components and behavioural measures were conducted in order to explore the relation between behavioural and ERP measures. In a first step, we calculated correlations (Pearson correlation coefficient) of means across conditions for behavioural indices and ERPs. In a second step, we investigated the relation between experimental effects in ERPs and behavioural measures.

Across conditions, P1 amplitudes were positively correlated to the accuracy in the testing session, *r* = .297, *p* < .05, possibly indexing that higher attention allocation during visual processing is related to higher overall accuracy. Concerning our experimental effect of associated valence in the form of the central negativity, correlation analyses showed that steeper learning slopes for negative words during the learning session were significantly related to increased negativity in the time range of 85 to 100 ms, *r* = .263, *p* < .05.

## Discussion

This study investigated whether early emotion-related ERP effects in response to letter strings would be based on associative learning of perceptual features or on fast access to associated valence information. In order to distinguish between these two options, we applied an associative learning paradigm where half of the letter strings were acquired in the visual domain, and the other half in the auditory domain with monetary gain, loss, or zero outcome.

In the test session, analyses of P1 amplitudes only showed trend-level results, pointing to an interaction between associated valence and learning modality. Exploratory tests at central electrode sites, however, indicated that early valence effects were limited to stimuli acquired in the visual domain, but were absent for stimuli learned in the acoustic domain; follow-up analyses at the P1 region of interest corroborated these findings. Therefore, our data provide tentative evidence that associative learning of a word’s shape might play a role in the emergence of valence effects in the visual cortex. Associative learning of perceptual features has previously been demonstrated for other symbolic stimuli (Rossi et al., 2017; Schacht et al., 2012), and also for pseudowords using an evaluative conditioning paradigm (Fritsch & Kuchinke, 2013; Kuchinke et al., 2015). However, by using a design where stimuli were acquired in two modalities, our study is the first to allow for a distinction between the influence of associated valence information and experience with a stimulus perceptual features. Furthermore, by careful counter-balancing of learning modalities and valence categories, we were able to avoid any perceptual differences between experimental conditions.

Early emotion effects in response to written words have previously been reported in a small number of studies (Bayer et al., 2012; Hofmann et al., 2009; Keuper et al., 2013, 2014; Kuchinke et al., 2014; Rellecke et al., 2011; Scott et al., 2009), but the boundary conditions of these effects remain unclear, especially since the majority of studies on emotional language processing failed to report these effects. Furthermore, emotion effects within such an early time range seem to contradict established reading models, which assume that lexico-semantic features are accessed only at around 200 ms after stimulus onset, while earlier time windows are indicative of orthographic analyses (for review, see Barber & Kutas, 2007). Our data suggest that early emotion effects do not necessarily contradict these models. Instead, they might be based on an additional mechanism, which might enable the system to quickly detect stimulus valence associated with a word’s perceptual features. Importantly, the notion of two distinguishable mechanisms underlying emotion effects in visual language processing does not imply their independence. As an example, interactions between associated and semantic valence might be at the core of findings like faster lexical access to emotional as compared to neutral words (Kissler & Herbert, 2013).

Reports of early emotion effects within 200 ms after stimulus onset in response to existing and associated/conditioned stimuli across stimulus domains are characterised by a marked heterogeneity concerning the direction of effects. While a number of studies report increased amplitudes for positive valence (Hammerschmidt et al., 2017; Schacht et al., 2012), others report increased amplitudes for negative valence (Kuchinke et al., 2015; Rossi et al., 2017). Our results are in accordance with the latter findings; we observed evidence for increased activation in the P1 time window for negative stimuli, mostly visible as an increased negativity at centro-parietal electrodes. The heterogeneity of early effects even in similar experimental paradigms suggests that this attention allocation is not hard-wired to a certain valence category, but might ultimately depend on specific task requirements and stimulus properties.

Concerning the effects reported in the present study, it has to be taken into consideration that the valence modulation in the P1 time range was observed on a group of central electrodes, in the form of an increased negative counterpart of the P1 for negative compared to neutral words acquired in the visual modality. Even though this effect was also evident at lateral P1 electrodes in exploratory post-tests, showing increased amplitudes for the (positive-going) P1 component, the experimental effects reported here are certainly small, especially considering the high number of subjects. As stated above, there is consensus that modulations within the first 200 ms are less stable and less reliably observed than later effects, likely due to their focal nature and their short duration. Furthermore, previous research suggests an influence of participant characteristics, such as trait anxiety, on the occurrence effects of associated valence (Rehbein et al., 2015). In order to corroborate the validity of our effects, we conducted exploratory brain-behaviour correlations, showing that the amplitudes of the central negativity from 85 to 100 for negative stimuli acquired in the visual domain was related to steeper learning slopes of those stimuli, indicating that faster acquisition was related to higher (negative) amplitudes.

Our finding of increased ERP amplitudes shortly after stimulus onset for negatively associated stimuli corroborates findings from (classical) conditioning studies. In these studies, stimuli are paired with aversive events like electric or acoustic shocks (e.g., Hintze et al., 2014; Rehbein et al., 2014), which elicit unconditioned reflex responses. Our study suggests that these reflex responses are not a prerequisite for the formation of stimulus associations that are able to impact the early phase of stimulus evaluation, but that associations can also be achieved when using secondary incentives, like money in the present study (for discussion, see Rossi et al., 2017).

In the present study, LPC amplitudes were not modulated by associated valence of letter strings. Previous findings were inconclusive with regard to occurrence and direction of LPC modulations (Fritsch & Kuchinke, 2013; Hammerschmidt et al., 2018, 2017; e.g., Schacht et al., 2012) and point towards a possible influence of task parameters (Rossi et al., 2017). As predicted, and in line with previous research, P300 amplitudes were increased for old letter strings compared to new distractors (Kuchinke et al., 2015; Rossi et al., 2017), reflecting the recollection of previously acquired stimuli (Rugg & Curran, 2007).

Despite the fact that early ERP effects were based on negative valence associations, behavioural data showed a clear advantage for positive associated valence. Participants made more correct classifications for positive stimuli than for negative and neutral stimuli during the learning session and achieved steeper learning slopes, indexing faster acquisition. In the test session, participants reacted faster and made fewer errors in response to positive letter strings. These results corroborate behavioural findings of associative learning studies using a highly similar design (Hammerschmidt et al., 2017; Rossi et al., 2017), showing a strong impact of reward on human perception and behaviour in service of optimizing goal-related behaviour (Anderson, 2013; Bourgeois, Chelazzi, & Vuilleumier, 2016; Navalpakkam, Koch, & Perona, 2009). Interestingly, the behavioural advantage for positive valence does not necessarily seem to translate to early valence effects in ERPs. While the advantage for positive valence corresponded to increased P1 effects for faces associated with positive valence in the study by Hammerschmidt and colleagues (Hammerschmidt et al., 2017), it was in other cases accompanied by increased amplitudes of the C1 in response to loss-associated stimuli (Rossi et al., 2017) and by increased amplitudes to loss-associated stimuli in the present study, despite almost identical learning procedures. One possible explanation relates to the stimuli used in these studies, since the first study used biologically meaningful stimuli, i.e. faces, while the latter ones used abstract symbols, thus possibly showing a differential sensitivity to associated valence across stimulus domains. Taken together, a number of studies using associative learning paradigms showed an advantage for reward-associated stimuli in behavioural parameters both during acquisition and when reward was no longer delivered, while the mechanisms underlying early modulations of visual stimulus processing remains to be fully understood.

## Conclusion

This study used an associative learning paradigm to associate meaningless letter strings with monetary gain, loss, or neutral outcome. After acquisition, stimuli associated with negative valence elicited increased ERP amplitudes around 100 ms after stimulus onset. Importantly, these effects were limited to letter strings acquired in the visual domain, but were absent for acoustically acquired stimuli. Thus, the present results indicate that associative learning of a word’s perceptual features might play a crucial role in the elicitation of emotion effects within 200 ms after word onset that was reported for existing words of emotional content.

## Footnotes

1 Please note that our learning procedure is purely a means of creating valence associations with letter strings, without the implication that this mimics actual word learning.

2 From a subset of 36 participants, we additionally collected data from an auditory testing session, which will not be presented here. For these participants, we recorded a lower number of trials in the visual testing session. Details are described in the procedure section.

